# Synergy Between TNFα and Proteostatic Stress Drives Cell Death and Guard Immunity

**DOI:** 10.1101/2024.06.12.598679

**Authors:** Matthew Sonnett, Edoardo Centofanti, Sanne Boersma, Max Brambach, Leon Peshkin, Roubina Tatavosian, Alon Oyler-Yaniv, Jennifer Oyler-Yaniv

## Abstract

The production and sensing of type I interferons (IFN-I) are critical for antiviral defense, yet most virus-infected cells do not produce IFN-I or upregulate IFN-stimulated genes. Using quantitative proteomics and global protein synthesis measurements, we show that productive viral infection globally down-regulates protein synthesis, restricting the IFN response. Guard immunity, which responds to disruptions in essential cellular processes, might compensate for the lack of IFN-I response by rapidly killing infected cells. However, non-pathological stressors can also disrupt proteostasis, making it unclear how cells decide to trigger guard immunity. We hypothesized that TNFα, produced by macrophages, provides a contextual signal allowing specificity. Using live-cell fluorescence microscopy and mathematical modeling, we showed that TNFα synergizes with the rapid decay of the anti-apoptotic protein c-FLIP to induce cell death and prevent viral spread. Our findings demonstrate that TNFα contextualizes proteostasis loss as non-sterile, enabling the activation of guard immunity to counteract viral infection.

## Introduction

The recognition of pathogen associated molecular patterns by germline-encoded pattern recognition receptors (PRR) is a central pillar of immunity ^1^. Once ligated, PRRs initiate a signaling cascade that culminates with the *de novo* production of type-I interferons α and β (IFN-I), which are secreted, sensed through extracellular receptors, and induce immune gene expression programs encompassing chemokines, cytokines, and restriction factors. The efficient production and sensing of IFN-I is critical for immunity to essentially all viruses, as individuals with inborn or acquired errors of the IFN-I signaling axis are more susceptible to a wide array of viral infections and severe disease ^2–5^.

Despite the importance of type I IFN for mounting a successful antiviral response, only a small fraction of virus-infected cells induce its expression or up-regulate IFN stimulated genes (ISG) ^6–12^. Several studies have demonstrated roles for both stochastic cell-intrinsic factors, and viral virulence factors in driving this fractional IFN response ^6,11–15^. However, the broad implication is that the vast majority of virus infected cells fail to mount an IFN-I response, which poses the question of how infected cells counteract or compensate for such immune impairment.

Viral infection, replication, and egress, are achieved by hijacking the host gene expression and trafficking machinery and antagonizing defense signaling. These actions profoundly disrupt normal cellular physiology. We therefore reasoned that a complementary branch of the immune system - guard immunity - can counteract impairment in the IFN-I response by sensing and responding to the same cellular disruptions that prevent IFN and ISG induction ^16,17^. Guard immunity was first described and primarily studied in plants, where plant resistance (*R*) genes guard the integrity of host proteins targeted by virulence factors ^18^. Disruption or modification of the host target activates the *R* protein, triggering an immune response. The plant hypersensitive response is a classic example and is characterized by rapid, localized cell death that limits pathogen spread and releases chemicals that warn neighbors. Recently, examples of guard immunity have been identified in vertebrates and mammals, and the definition has expanded to include sensing of disruptions to essential cellular processes, such protein synthesis or ion flux ^16,17^.

A key open question in understanding guard immunity is how specificity can be achieved. Many environmental stressors, such as transient nutrient deprivation or exposure to ultraviolet (UV) radiation, perturb essential cellular processes but pose no broader danger to the organism. It is not known how a cell would differentiate between a *sterile* perturbation, and a perturbation brought about by a pathogen. Owing to their ubiquitous distribution in host tissues and constitutively high expression of PRRs ^19^, macrophages and other innate immune sentinels are the first cells to sense and respond to incoming pathogens. When macrophages sense the presence of pathogens, they rapidly secrete pro-inflammatory cytokines including interleukins (IL)-1, 6, 12, type I and II interferons, and Tumor Necrosis Factor (TNFα). Our previous work demonstrated that TNFα dramatically hastens cell death in virus-infected cells ^20^, leading us to hypothesize that, in vertebrates, cytokines might contextualize viral disruptions to essential cellular processes, and thereby provide a licensing signal for the activation of guard immunity.

To test this hypothesis, we examined the role of TNFα in the relationship between viral virulence and guard immunity during Herpes Simplex Virus-1 (HSV-1) infection. Using quantitative proteomics and global measurements of protein synthesis, we found that productive virus infection leads to significant, global, and progressive down-regulation of protein synthesis and failure to initiate an antiviral or IFN response. Combining live-cell fluorescence microscopy with mathematical modeling, we demonstrated that TNFα signaling synergizes with the rapid decay of the anti-apoptotic protein c-FLIP—identified here as a novel guard factor for proteostasis—to kill infected cells and prevent viral spread. Importantly, while other sterile environmental stressors can cause similar translational suppression, cell viability remains unaffected without TNFα. Thus, TNFα contextualizes the loss of proteostasis as ‘non-sterile,’ enabling specificity in cellular guard immunity.

## Results

### Infection with virulent viruses leads to genome-wide protein down-regulation

Attenuated viruses generally activate the type I IFN response more effectively than their virulent counterparts ^21^. We therefore asked how progressively-attenuated HSV-1 strains would impact host antiviral protein expression. To this end, we used tandem mass tags (TMT) ^22^ to perform 18-plex, quantitative, and time-resolved mass spectrometry on mouse fibroblasts infected with a panel of HSV-1 mutant strains (Fig 1A). We infected cells with an MOI 10 of the HSV-1 mutant strains d109, 8ΔGFP, dl38, or full-length virus all on the parental KOS strain background. These strains are variably-attenuated as they lack the ability to activate viral gene expression (d109 ^23^), replicate the genome (8ΔGFP ^24^), or assemble viral capsids (dl38) (Fig 1B). Henceforth, these strains will be referred to as strongly-attenuated (d109), moderately-attenuated (8ΔGFP and dl38), and wild type. This MOI resulted in >90% of the cells becoming infected from the initial viral inoculum, as measured by immunofluorescence staining of the immediate-early protein ICP4 (Fig S1A). Infection with the strongly-attenuated d109 strain could not be confirmed in this way as it does not express any viral genes.

**Figure 1:**
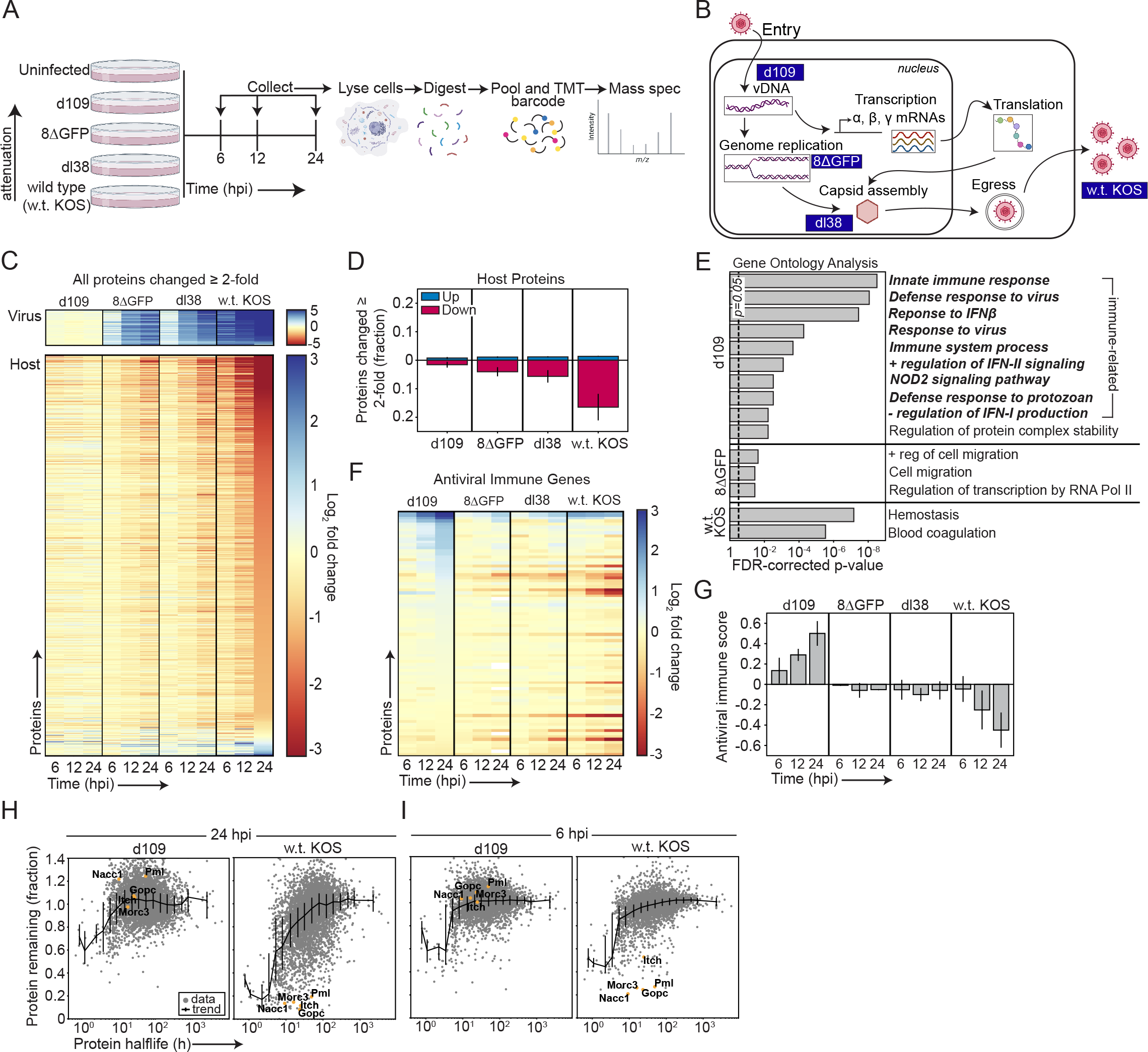
*Viral infection leads to genome-wide protein down-regulation and immune impairment.* **(A)** Cartoon diagram showing the experimental layout for proteomics experiments. 3T3 fibroblasts are infected with an MOI 10 of each of the various HSV-1 strains. At the indicated time points, cells are harvested, and lysed. Then proteins are digested and labeled with TMT barcodes before LC/MS/MS. **(B)** Cartoon diagram showing HSV-1 viral life cycle and where each viral mutant used in this study is disrupted within the life cycle. **(C)** Heatmap showing all viral and host proteins that change by greater than or equal to 2-fold relative to the control (uninfected) condition. All proteins are ranked according to the 24 h time point of the wild type virus-infected cells. **(D)** Quantification of the total host proteins that are increased or decreased by greater than or equal to 2-fold relative to the control condition. The data is presented as the mean ± s.d. of two biological replicates. **(E)** Gene Ontology analysis of the top 200 increased proteins for cells infected with each of the different viral strains. A maximum of the top 10 statistically-significantly enriched pathways are shown for each condition. Note that there were no significantly enriched pathways detected for cells infected with the dl38 mutant. Immune-related pathways are emphasized in bold and italics. **(F)** Heatmap showing protein expression for our compiled list of antiviral immune genes. **(G)** Quantification of the antiviral immune score for cells infected with each of our viral mutants. To compute the score, we took the mean expression of all proteins on our antiviral immune list ± s.e.m. for each protein. **(H-I)** Quantification of the fraction of each protein compared to the uninfected control versus its published half-life for 24 hpi (H) or 6 hpi (I). The black trendline shows the average expression ± the 25 and 75% confidence intervals for binned proteins. Known virus-targeted proteins are annotated and shown in orange.

We quantified 11,015 total host proteins and 71 viral proteins, or 98% of known HSV-1 proteins ^25,26^. Viral proteins were dramatically up-regulated in infected cells, with exception of cells infected with the strongly-attenuated strain (Fig 1C). The moderately-attenuated strains exhibited reduced viral protein expression relative to the wild type strain. Of host proteins, between 3-20% changed by more than 2 fold, and most proteins were reduced in expression (Fig 1C-D). The degree of protein down-regulation scaled inversely with the extent of viral attenuation: more attenuated strains caused less down-regulation of host proteins.

These results are consistent with previous work that demonstrated infection with replicative HSV-1 strains (KOS and 17), caused significant down-regulation in protein expression in human immortalized keratinocytes and primary fibroblasts ^27–29^. In addition, previous studies have demonstrated a similar predominance of down- versus up-regulated proteins during lytic infection with various other viruses including vaccinia virus, HIV, Epstein-Barr, and human cytomegalovirus (HCMV) ^30–33^. This, in combination with our and others studies of HSV-1 ^27–29^, indicate that genome-wide protein down-regulation is a common feature of many viral infections and is dependent on virulence. Our observations also highlight the utility of proteomics to study host and viral gene expression during viral infection. Our data suggests that mRNA will correlate poorly with protein, especially for induced genes, as has been demonstrated previously by ribosome footprinting of host and viral gene expression during SARS-CoV-2 infection ^34^.

### Antiviral proteins are only induced in cells infected with a strongly-attenuated virus

We next performed Gene Ontology (GO) analysis to identify biological pathways enriched among the top 200 up-regulated host proteins for each of the virus strains ^35,36^. While there was no enrichment for overtly immune-related pathways in cells infected with the wild type or moderately-attenuated strains, infection with the strongly-attenuated strain led to significant enrichment of proteins involved in antiviral innate immunity and the type I IFN response (Fig 1E). We used GO annotations and literature review to compile an antiviral immune signature of proteins involved in the host innate response to viral infection. As expected, many of these proteins are up-regulated in cells infected with the strongly-attenuated mutant, yet progressively down-regulated in cells infected with the more virulent strains (Fig 1F). We did not observe a similar pattern of up-regulation for two sets of proteins not directly involved with innate immunity, yet did observe down-regulation (Fig S1B-C). To quantify this, we computed an antiviral immune expression score, by taking the mean protein expression for all of the proteins on the antiviral gene list. This shows that cells infected with the strongly-attenuated mutant increase expression of antiviral genes over time (Fig 1G). By contrast, cells infected with the moderately-attenuated strains fail to induce expression of antiviral proteins, and the wild type strain leads to significant and progressive reduction in antiviral protein expression.

To test whether the pattern of up- and down- regulation was specific to antiviral immune genes, or reflected a global pattern affecting the entire proteome, we compared the distribution of protein fold expression changes to 10,000 randomly selected length-matched gene lists. To measure distribution similarity, we calculated the average Kolmogorov-Smirnov statistic for different gene sets. We found that the pattern of antiviral protein expression in cells infected with any but the strongly-attenuated virus is statistically indistinguishable from that of a random gene set (Fig S1D-E). In contrast, the increase in antiviral protein levels in cells infected with the strongly-attenuated strain is statistically distinct from random, indicating virus-specific regulation. Corroborating these results, an expression score for an unrelated gene set shows absence of protein up-regulation, but comparable attenuation-dependent down-regulation (Fig S1F). These results argue that while immune protein up-regulation by cells infected with the strongly-attenuated strain is virus-specific, the antiviral down-regulation observed during wild type infection aligns with a model of global, non-specific protein decay.

The strongly-attenuated mutant can enter the cell and deposit its genome, but cannot transcribe viral genes ^23^. It therefore creates an infection similar to defective or abortive viral replication, which can be caused by host factors and defective viral genomes ^9,37–39^. Abortive infection occurs when cells become infected and express immediate-early genes, yet never progress to late gene expression and progeny production ^9^. Previous single cell studies of the transcriptome demonstrate that only rare subpopulations of infected or abortively-infected cells are capable of initiating a type-I IFN or antiviral response ^7–9,40^. Our data corroborate these findings at the bulk protein level, and further confirm that only cells that control the virus early in infection can initiate an innate immune response. Taken together, our data show that productive viral infection not only impairs the induction of antiviral genes, but also significantly decreases their expression.

### Productive viral infection causes non-specific protein decay characteristic of halted translation

If the global protein decay that we observe is due to translational inhibition, we reasoned that protein degradation would most significantly impact short-lived proteins. To examine this, we plotted the fraction of remaining protein at 24 hpi versus the published half-lives ^41^ for each protein in the same cell line. We compared between cells infected with the d109 mutant, which causes minimal protein decay, and the wild type strain, where decay is significant. This analysis supports our hypothesis that decay is due to translation inhibition by showing that short-lived proteins are reduced to much greater extent than proteins with long half lives (Fig 1H).

Herpesviruses - including HSV-1, -2, and HCMV can specifically target certain host proteins including Gopc, Itch, Morc3, Pml, and others, for degradation ^27,30,42–45^. Many of these proteins are associated with Pml-nuclear bodies, which are a component of cell intrinsic immunity and hence the cells’ first line of defense ^46^. Indeed, each of these proteins was significantly down-regulated by 6 hpi and remained so until at least 24 hours, during infection with the wild type strain, but not the d109 mutant. (Fig 1H-I). Hence, although some proteins are directly targeted for destruction by the virus, the broad immune impairment we observe in virus-infected cells arises as a global, indirect consequence of translational inhibition, as has been suggested for other viruses ^34,47^.

### Infected cells reduce protein synthesis commensurate with viral attenuation levels

Our data suggests that HSV-1 infection halts host gene expression (Fig 1). Previous work has shown that, although total mRNA is reduced during HSV-1 infection, transcription remains largely constant ^48^. This led us to hypothesize that the global protein down-regulation we observed across the different HSV-1 mutant strains is due to progressive suppression in translation. We used puromycin incorporation ^49^ to test this hypothesis and quantify nascent protein synthesis after infection with MOI 10 of each HSV-1 strain (Fig 2A). Puromycin mimics tyrosyl-tRNAs and is incorporated into the elongating peptide. After fixation and permeabilization, puromycilated peptides are detected using immunofluorescence, and the fluorescence intensity can be used as a read-out for *de novo* protein synthesis.

**Figure 2:**
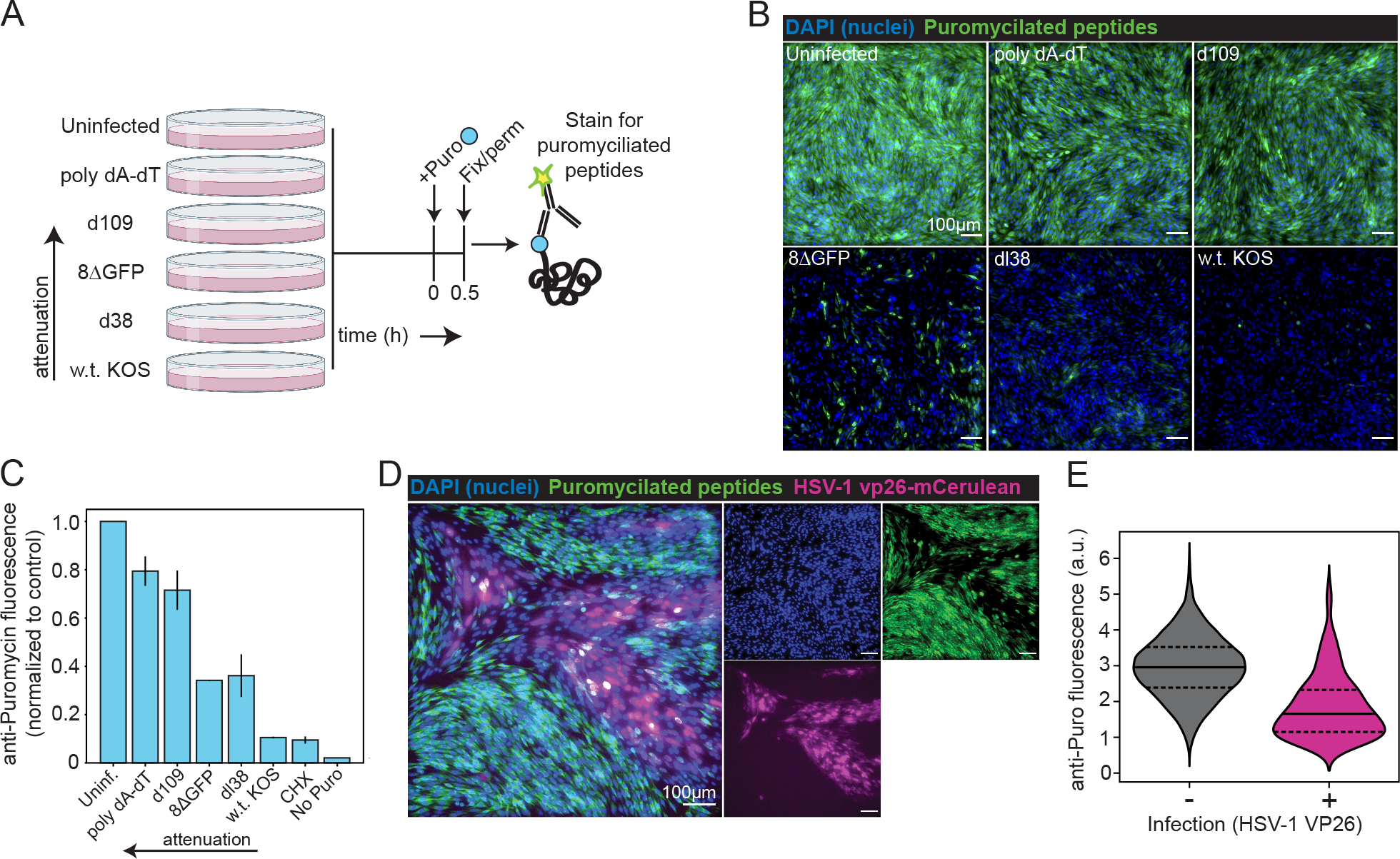
*Infected cells reduce protein synthesis commensurate with viral attenuation levels.* **(A)** Cartoon diagram depicting the experimental layout for puromycin incorporation assays. Cells are first infected with various viral strains, then pulsed for 30 min with 2 μM puromycin before fixing, permeabilizing, and staining for puromycilated peptides. **(B)** Representative images of puromycin staining for cells infected overnight with MOI 10 of each of the virus strains or transfected with 200 ng/well of poly dA-dT. **(C)** Quantification of average puromycin fluorescence per cell for each of the different conditions. Bars show the mean ± s.d. of multiple biological replicates. **(D)** Representative images of puromycin staining for cells infected overnight with MOI 3 of full length, fluorescent-tagged HSV-1. **(E)** Violin plot of anti-puromycin fluorescence for VP26-mCerulean+ (infected) or - (uninfected) cells. Inner lines show the median ± quartiles of the data.

We observed potent reduction in translation that is commensurate with the extent of viral attenuation (Fig 2B-C). Translational interference is a common consequence of viral infection, and has previously been observed during infection with HSV-1 ^50^, coronaviruses ^34,47,51^, and other viruses ^52^. We also observed marked reduction in protein synthesis during infection with Influenza A virus (IAV/WSN/1933) in both mouse (Fig S3A-B) and human (Fig S3C-D) cells, and for human cells infected with HSV-1 (Fig S3E). The mechanism responsible for translational suppression is likely multi-faceted and originates from both virus and host factors as transfection of fibroblasts with host shutoff genes from HSV-1 or SARS-CoV-2 (*Vhs* or *Nsp1*, respectively) causes only a partial reduction in protein synthesis relative to full length HSV-1 (Fig S3F). Finally, we checked whether translational inhibition was restricted to infected cells by infecting with a lower MOI of 3, resulting in patches of infected cells. Only productively-infected cells exhibit translational inhibition, consistent with what others have shown (Fig 2D-E) ^50^. Taken together, these data demonstrate that infection with variably-attenuated HSV-1 strains results in graded suppression in protein synthesis. More broadly, suppression of translation appears common as it occurs during other viral infections and in human cells.

### TNFα synergizes with translational inhibition to induce cell death during virus-infection

Viral infection leads to translational repression, which hamstrings the production of antiviral proteins. This raises the question of whether and how translationally-repressed cells are able to restrict viral spread. Programmed cell death is a powerful cell-autonomous antiviral mechanism: If it occurs rapidly after the onset of infection, it restricts the virus from spreading ^53^. Notably, cell death induction requires pre-made proteins and therefore does not depend on *de novo* protein synthesis. Programmed cell death may therefore serve as an alternative antiviral mechanism that can operate effectively during inhibition of protein synthesis.

We previously showed that viral infection sensitizes cells to the cytotoxic cytokine Tumor Necrosis Factor-α (TNFα), which restricts viral spread in a cell death dependent manner ^20^. Others have shown that inhibiting translation sensitizes cells to the related cytotoxic cytokine, TRAIL ^54,55^. We therefore asked whether the increased sensitivity to TNFα observed during viral infection ^20^ is due specifically to synergy of TNFα with translational inhibition. To test this hypothesis, we first inhibited protein synthesis orthogonally using an established protocol employing translation inhibitors cycloheximide (CHX) and anisomycin (ANM), and quantified cell death synergy when combined with TNFα ^56^.

We co-treated cells with a low dose of TNFα and CHX or ANM, and imaged cell death over time via incorporation of a cell impermeant dye (Fig 3A-D). Translation inhibition alone does not kill cells over the course of 18 h, yet combination with TNFα synergizes potently to induce rapid and nearly complete death of the entire culture (Fig 3C-D). Previous work has shown that turnover of short-lived pro-survival proteins couples the cessation of translation to apoptosis ^57^. Consistent with this model, addition of a proteasome inhibitor cocktail (Prot_I_) delayed cell death for TNFα-treated, translation-inhibited cells (Fig 3C-D). This experiment suggests that translation inhibition causes proteasome-mediated degradation of proteins that antagonize extrinsic apoptosis. To quantify cell death time course trajectories and enable more straightforward comparison between different experiments, we measured death rates by fitting each curve to an exponential function as we described previously (Fig 3E-F) ^20^.

**Figure 3:**
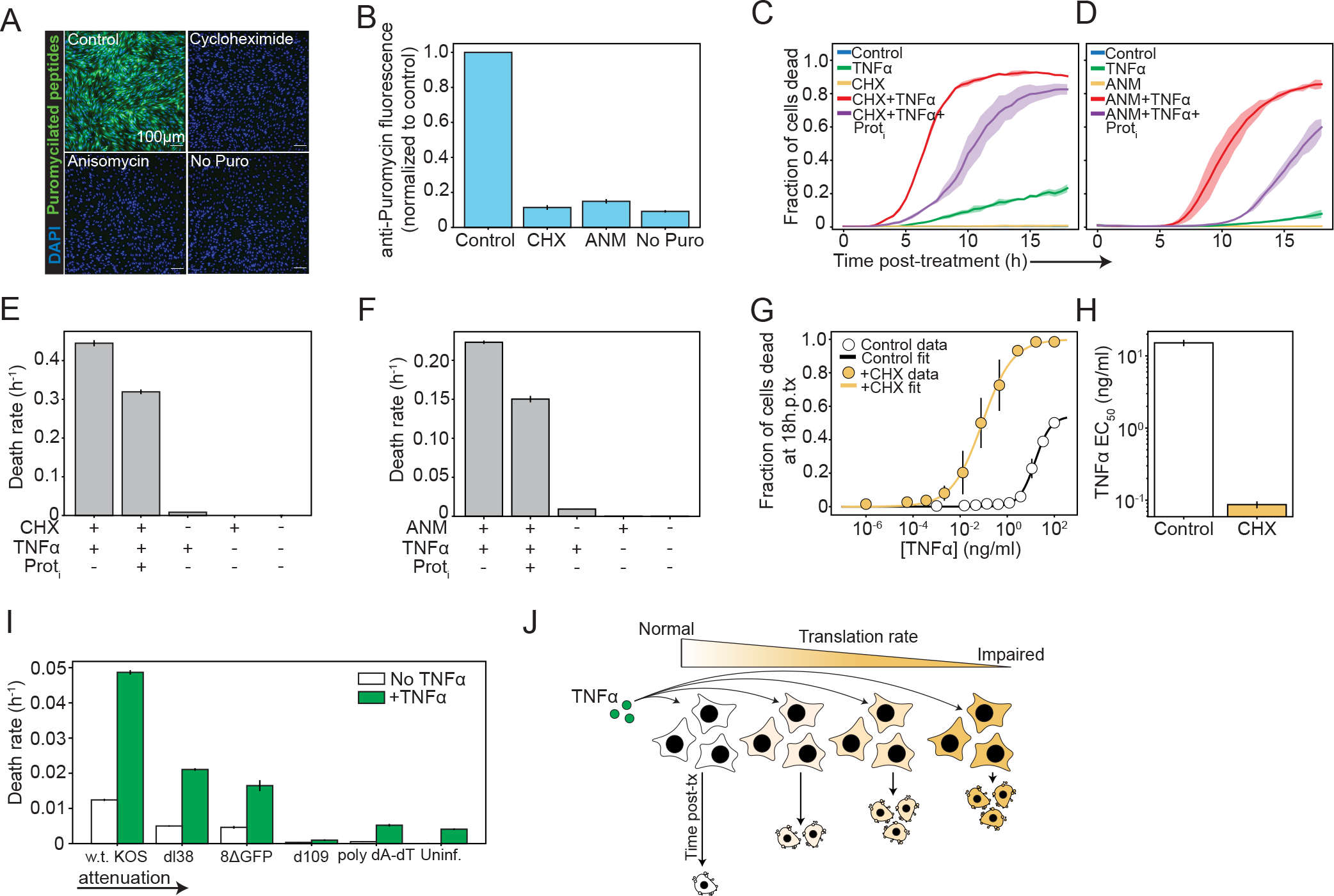
*Perturbing protein synthesis dramatically sensitizes cells to TNFα.* **(A)** Representative images of puromycin staining for fibroblasts pre-treated for 20 min with 20 μM CHX or 5 μM ANM prior to starting the puromycin pulse. **(B)** Quantification of the average cellular puromycin fluorescence intensity from A. **(C)** Quantification of cell death over time (based on Sytox green incorporation) for cells treated with either 20 μM CHX or 5 ng/ml TNFα, CHX with TNFα, or CHX with TNFα and our proteasome inhibitor cocktail (Prot_I_ - 2 μM bortezomib/2.5 μM carfilzomib). **(D)** Quantification of cell death over time for cells treated with either 5 μM ANM or TNFα, ANM with TNFα, or ANM with TNFα and Prot_I_. **(E-F)** Quantification of death rates fit from C and D. Death rates were fit by converting cell death to log space and linearly fitting the exponential component, as we did previously ^20^. Error bars show the standard error of the fit. **(G)** Quantification of cell death at 18 hp-tx (based on Sytox green incorporation) for cells treated with a dose titration of TNFα with or without concurrent treatment with 20 μM CHX. Curves were fit with a Hill function with coefficient 1. **(H)** TNFα EC_50_ values computed from H. Error bars show the standard error of the fit. **(I)** Quantification of death rates for cells infected with MOI 10 of each virus strain with or without 5 ng/ml TNFα. The error bars show the standard error of the fit. **(J)** Cartoon diagram showing the working model: Translational inhibition tunes the cellular sensitivity to TNFα. For fig 3, unless otherwise stated, the error bars show the mean ± s.d. of multiple biological replicates.

We noticed that proteasome inhibition delays cell death, but does not block it. Previous work has demonstrated that proteasome inhibition also sensitizes cells to TRAIL-mediated death by preventing turnover of enzymatically active (cleaved) Caspase 8 ^58^. Consistent with this, we showed that proteasome inhibition also sensitizes cells to TNFα-mediated cell death, yet not to the same extent as translational inhibition (Fig S3A-B).

We measured the increase in TNFα sensitivity conferred by translational inhibition by treating CHX or vehicle-treated cells with a dose titration of TNFα and quantifying total cell death 18 hours later (Fig 3G). We fit these curves with a Hill function, which reveals that translational inhibition makes cells ∼100-fold more sensitive to TNFα (Fig 3I).

If our model of synergy between translational inhibition and TNFα treatment is correct, we expect that graded translational inhibition caused by attenuated virus strains will translate into graded synergy with TNFα. We tested this hypothesis by infecting cells with an MOI 10 of each virus strain, treating with low dose TNFα, and quantifying cell death over time. Consistent with our past work ^20^, neither virus was very cytotoxic alone, yet combination with TNFα synergistically enhanced cell death in a manner that scaled inversely with viral attenuation (Fig 3I, and S3C). We noticed that the strongly-attenuated viral mutant actually protected cells against TNFα-mediated cell death compared to the uninfected control. This suggests that the protein expression changes induced by the strongly-attenuated mutant confer a pro-survival cell state. Indeed, previous work demonstrates that defective viral genomes (DVG), which are conceptually similar to a strongly-attenuated virus, promote cell survival in response to TNFα, facilitating long-term persistence of paramyxoviruses ^59^. These data demonstrate the complex effects that DVGs or abortive infections can have on host immunity and viral fitness: On the one hand, they enable the host to efficiently activate innate immune protein expression, yet may also generate heterogeneous cell states that facilitate viral persistence.

Finally we confirmed that these observations are reproducible in response to other cytotoxic ligands (Fas), and in other cells (human A549) (Fig S3D-G). Taken together, our data demonstrate that graded inhibition of protein synthesis caused by viral infection or translational inhibitors is quantitatively converted into graded sensitivity to cytotoxic cytokines (Fig 3J). The fact that proteasome inhibition delays cell death implies that turnover of short-lived pro-survival proteins mediates this effect.

### Synergy between TNFα and translation inhibition accelerates cell death to restrict viral spread

We previously showed that TNFα dose-dependently slows the spread of virus ^20^. By shortening the time interval between the onset of infection and death, the virus cannot complete its life cycle and does not spread (Fig 4A). Demonstrating the efficiency of this simple antiviral defense, we showed, using epidemiological simulations, that sufficiently-rapid cell death enacted early in infection can eliminate the virus from the population altogether ^20^. We therefore sought to test whether TNFα inhibition of viral spread requires intact cell death signaling.

**Figure 4:**
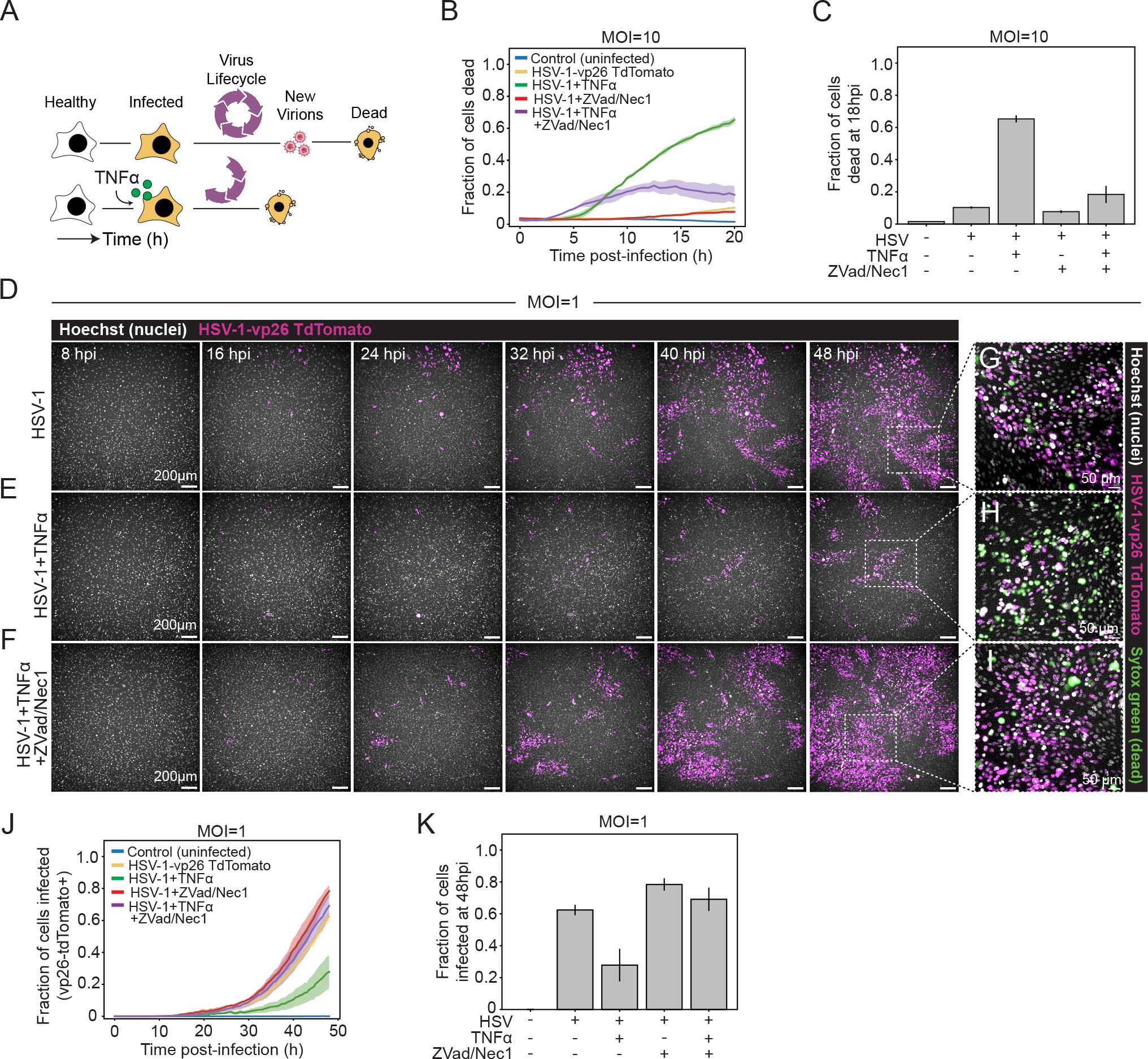
*Accelerated cell death restricts viral spread.* **(A)** Cartoon diagram showing the principle. TNFα accelerates cell death after viral infection, which prevents completion of the viral life cycle and production of new infectious virions ^20^. **(B)** Quantification of cell death over time for cells infected with MOI 10 of full length HSV-1 alone, or HSV-1 with TNFα, HSV-1 with 30 μM each of ZVad and Nec1, or HSV-1,TNFα and ZVad/Nec1. **(C)** Quantification of cell death with the indicated conditions at 18hpi. **(D-F)** Images showing viral spread for cells left untreated (D), treated with TNFα (E), or treated with TNFα and Z Vad/Nec1 (F). **(G-I)** Close-up images showing cell death from within insets of larger images for each condition. **(J)** Quantification of infected cells over time (based on expression of HSV-1 VP26-tdTomato) after infection with MOI 1 for the indicated conditions. **(K)** Quantification of infected cells for each of the indicated conditions at 48hpi. For all of fig 4, plots show the mean ± s.d. of multiple biological replicates.

To do so, we first quantified the effect of pharmacological inhibition of cell death signaling on cell death during viral infection. Cells were infected with an MOI 10 of HSV-1, and treated with a low dose of TNFα with or without addition of the pan-Caspase inhibitor ZVad and RIP kinase inhibitor Necrostatin-1 (Nec1) to inhibit cell death signaling. The combination of ZVad and Nec1 strongly reduced programmed cell death in TNFα-treated, virus infected cells (Fig 4B-C). This shows that inhibiting programmed death signaling pathways blocks the execution of cell death provoked by TNFα in virus-infected cells.

We hypothesized that the protective effect of TNFα on restriction of viral spread is mediated by execution of programmed cell death. If true, blocking cell death will rescue viral spread and abrogate TNFα’s protective effect. To test this, we treated cells with a low MOI (1) of fluorescent-tagged HSV-1 ^60^ and measured viral spread over time using microscopy. These images, along with their systematic quantification, show that while TNFα has no effect on the number of initially infected cells, it potently restricts viral spread, resulting in only small patches of infection compared to untreated cells (Fig 4D-E, J-K). Inhibiting programmed death signaling using ZVad and Nec1, completely abrogates the protective effect of TNFα, indicating that the protective effect of TNFα depends on cell death (Fig 4F, J-K). Examination of patches of infected cells shows a high density of dead cells within infection clusters in TNFα-treated cultures (Fig 4G-I). Taken together, these experiments show that TNFα, which is normally released by activated macrophages, acts as a danger cue by guiding viral infected cells to die, and thus limiting viral spread.

### c-FLIP guards for perturbations in proteostasis

We’ve shown that viruses perturb protein synthesis, leading to decay of short-lived proteins and sensitization to TNFα that triggers proteasome-dependent cell death. This implies that there are fast-degrading pro-survival proteins which guard for viral perturbations in protein synthesis. There are several guard candidate proteins embedded in the apoptotic signaling network that negatively-regulate extrinsic apoptosis: Bcl2 family members, which antagonize pore-forming enzymes in the mitochondrial membrane; Xiap, which inhibits Caspase 3; and c-FLIP, which inhibits Caspase 8 (Fig S4A) ^61^.

We reasoned that if any of these candidate proteins guard for proteostasis, then inhibiting or down-regulating them and co-treating with TNFα will phenocopy the synergistic effect of translational inhibition combined with TNFα. Previous work identified a role for Bcl2 members BclXL and Mcl1 in mediating programmed cell death during viral infection ^57,62^. Hence, we first tested whether inhibiting Bcl2 family members using a pan-Bcl2 inhibitor (Obatoclax) or specific Mcl1 inhibitor (A1210477) sensitized cells to TNFα. However, neither compound induced cell death in combination with TNFα (Fig S4B-C). Xiap inhibition - using the SMAC mimetic LCL-161 - did accelerate the onset of cell death when combined with TNFα, yet only killed half as many cells as translational inhibition (Fig S4D).

However, siRNA knockdown of c-FLIP (Cflar) phenocopied the effect of translational inhibition using CHX or ANM, by synergizing with TNFα to induce cell death (Fig 5A-B and S4E). We reasoned that the protective effect of proteasome inhibition (Fig. 3 C-F) would be nullified in our Cflar knockdown cells, as siRNA knockdown is proteasome independent (Fig 5A). By contrast, CHX-mediated protein decay depends on the proteasome. Indeed, death induction was insensitive to Prot_I_ in Cflar siRNA-treated cells but not in the scramble, CHX-treated control (Fig 5C and S4E). Similar to translation inhibition using CHX, Cflar knockdown decreased the TNFα EC_50_ by ∼100 fold (Fig 5D-E and 3G-H). Consistent with this result, our proteomics data reveals graded loss of c-FLIP during infection that scales inversely with the extent of viral attenuation, which is confirmed by western blotting (Fig 5F, S4F). We therefore propose that c-FLIP (Cflar) performs a guard function for protein synthesis: Loss of translation leads to rapid decay, which lowers the threshold for TNFα-mediated cell death by nearly 100-fold.

**Figure 5:**
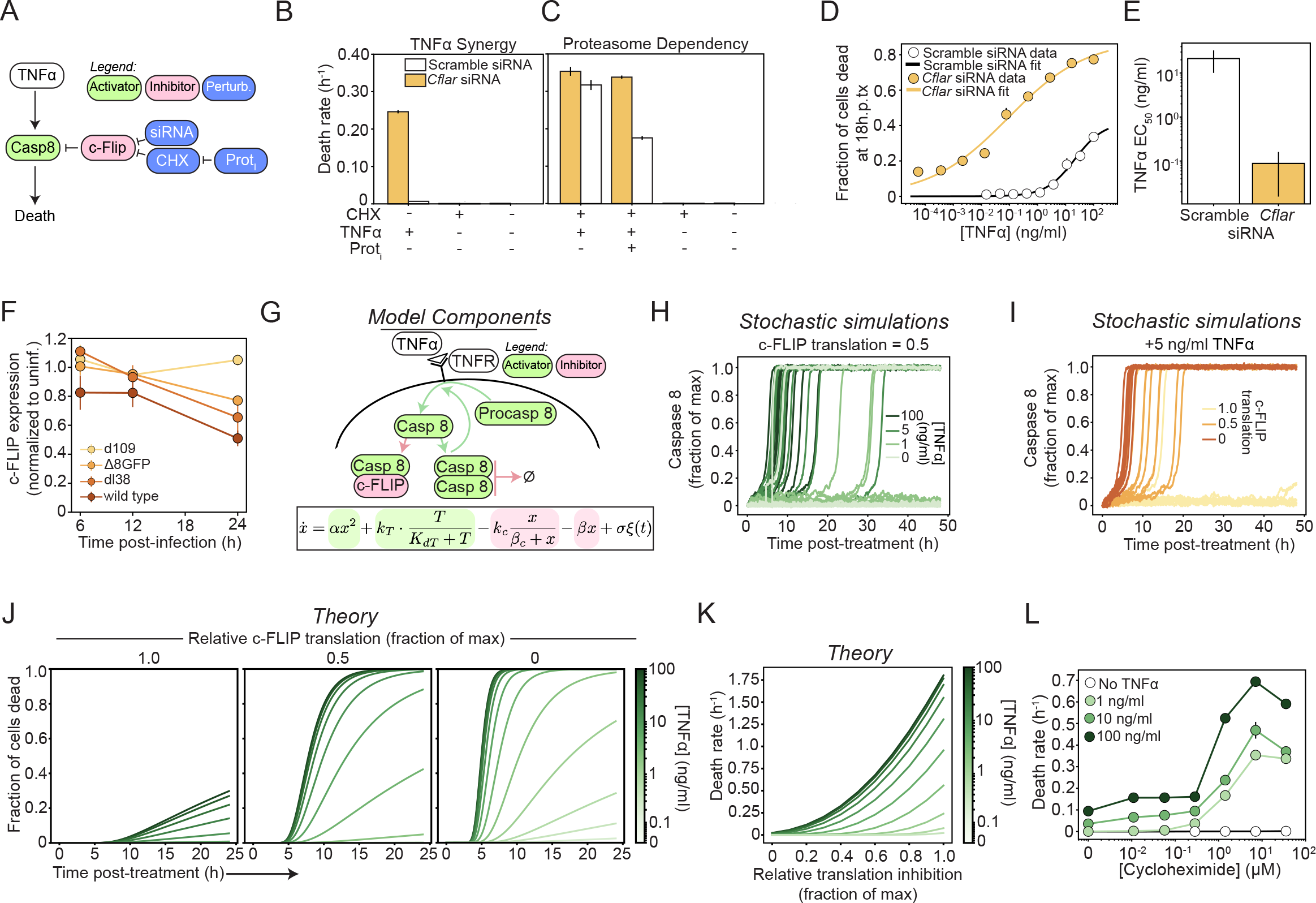
*c-FLIP guards for perturbations in protein synthesis.* **(A)** Cartoon diagram of the key relationships between molecular components perturbed in 5B-C. **(B-C)** Quantification of death rates for *Cflar* or control siRNA treatments. Error bars show the standard error of the fit. **(D)** Quantification of dead cells for control siRNA or *Cflar* siRNA 18hp-tx with a dose titration of TNFα. Each data point shows the mean ± s.d. of multiple biological replicates. Data were fit with a Hill function. **(E)** TNFα EC_50_ values computed from C. Error bars show the standard error of the fit. **(F)** Quantification of c-FLIP over time from proteomics experiments for cells infected with each of the viral mutants. Data show the mean ± s.d. of biological replicates normalized to the control (uninfected condition) at each relevant time point. **(G)** Cartoon diagram of the mathematical model. Our model incorporates TNFα triggered Procaspase-8 cleavage, autocatalysis of Procaspase-8 by Caspase-8 dimers, inhibition of Caspase-8 by free c-FLIP, natural Caspase-8 degradation, and stochastic biochemical noise. **(H)** Individual trajectories for stochastic simulations of Caspase 8 cleavage for three different TNFα concentrations and a c-FLIP translation rate of 0.5 of the maximum. **(I)** Individual trajectories for stochastic simulations of Caspase 8 cleavage for three different c-FLIP translation rates and a fixed low concentration of TNFα. **(J)** Model predictions for the proportion of cells killed by treatment with different concentrations of TNFα and three different rates for c-FLIP translation. **(K)** Model predictions of the population death rates for a spectrum of different TNFα concentrations and c-FLIP translation rates. **(L)** Quantification of the death rates for cells treated with dose titrations of cycloheximide and the indicated concentrations of TNFα. Error bars show the standard error of the fits.

### A noise induced stochastic switch drives the timing of death decisions

We were intrigued by the extent to which relatively modest decreases in c-FLIP protein expression synergize with TNFα to induce cell death. We therefore turned to mathematical modeling of Caspase 8 activation to quantitatively probe the characteristics of the apoptotic signaling network that permit such cell death dynamics. Signaling in response to TNFα scales with the dose of ligand ^63,64^, yet there is a high degree of variability both between cells and for a given cell over time. Variability in signaling and in downstream cell death decisions is driven by differences in protein expression, along with stochastic biochemical noise ^54,58^, calling for the use of a stochastic differential equation model (Fig 5G). Downstream of receptor signaling, Procaspase-8 is cleaved into active Caspase-8 in proportion to the level of receptor activation ^55^. Activated Caspase-8 molecules must dimerize, before further inducing Procaspase-8 cleavage in an autocatalytic loop ^65^. c-FLIP acts as a classic competitive inhibitor of apoptosis by binding to Caspase-8, preventing dimerization and blocking its proteolytic activity ^58,66^. In addition, Caspase-8 degrades naturally via the proteasome ^58,67^.

Conceptually, the rate of c-FLIP translation dictates the ability of a cell to counteract activated Caspase-8. If Caspase-8 levels saturate all available c-FLIP, positive feedback will dominate, levels of Caspase-8 will rise rapidly, and the cell will die. Similar saturated removal models have been employed to explain the sharp increase in death rates during aging ^68^. Biochemical noise plays a crucial role in such systems as it can stochastically tip the levels of Caspase-8 above the saturation limit, triggering cell death. TNFα signaling increases the baseline level of Caspase-8 cleavage, such that a smaller - and therefore more likely - fluctuation will induce death (Fig 5H). Reduced c-FLIP translation lowers the threshold for Caspase-8 positive feedback activation, resulting in faster and less variable timing of cell death (Fig 5I).

After an initial lag caused by transition of the signaling pathway away from homeostasis, which we observe experimentally (Fig. 3, S3), different levels of TNFα and c-FLIP translation cause an exponential increase in the fraction of dead cells (Fig 5J-K). Our model predicts that in the presence of TNFα, a linear decrease in protein translation will cause a superlinear increase in cell death. Alone, low to moderate concentrations of TNFα or c-FLIP reduction are insufficient to cause much - if any - cell death. However, TNFα strongly synergizes with decreases in c-FLIP translation, driving rapid cell death. To test these predictions, we treated cells with a dose-titration of CHX, which caused a linear decrease in protein synthesis (Fig S4G) and measured death rates across different concentrations of TNFα. As predicted by the model, we observed a TNFα-dependent superlinear relationship between the degree of translation inhibition and the rate of cell death (Fig 5L). Furthermore, and in agreement with our proteomics measurements (Fig. 5F), even a relatively modest 50% drop in c-FLIP levels was sufficient to hasten the rate of cell death by orders of magnitude.

Hence, our model suggests, and experiments reinforce, an elegant molecular logic whereby c-FLIP levels regulate the probability that Caspase-8 positive feedback is activated in response to TNFα. The regulation of Caspase-8 positive feedback by stochastic biochemical noise enables a remarkably wide range of death rates to be achieved depending on the extent of translational suppression and TNFα stimulation.

### Coupling cell death to TNFα safeguards cell viability during sterile perturbations to protein synthesis

Many sterile stressors cause the cell to transiently inhibit translation, but pose no broader threat to the organism. Such instances are largely aimed at cellular recovery and promote cell survival and tissue health. However, the aberrant release of TNFα has deleterious effects in many disease contexts. Our data suggests that the release of TNFα in contexts of sterile proteostatic stress may inappropriately trigger guard immunity and cell death, despite the absence of a pathogen. To test this hypothesis, we exposed cells to several sterile stressors either alone, or in combination with TNFα, and quantified cell death over time.

The integrated stress response (ISR) responds to a broad array of different stressors by triggering one of four kinases (GCN2, PERK, HRI, or PKR), which phosphorylate the downstream eIF2α, halting translation of most mRNAs. To activate the ISR, we used several different approaches: UV irradiation and Thapsigargian, which induce ER stress and the kinase PERK; and Halofuginone, which inhibits aminoacyl tRNA synthetases to stimulate the amino acid starvation response and the kinase GCN2 (Fig 6A) ^69,70^. Consistent with the established consequences of each treatment, we observed significant inhibition of protein synthesis (Fig 6B-E). Further, although neither treatment caused much cytotoxicity alone, co-treatment with TNFα strongly synergized to induce cell death, similar to viral infection or translation inhibitors (Fig 6F-H and Fig 3C-F, and I, and S5A-C). These data indicate that synergy in cell death induced by the combination of proteostatic stress and TNFα is a general mechanism, because it is independent of the cause for translational arrest (Fig 6I). Because TNFα is released by activated macrophages in the context of pathogen infection, we conclude that it contextualizes loss of proteostasis as ‘non-sterile,’ thereby enabling specificity in guard immunity. Finally, conditioning the synergy in cell death on an immune-derived cue safeguards cell viability in contexts of sterile proteostatic stress.

**Figure 6:**
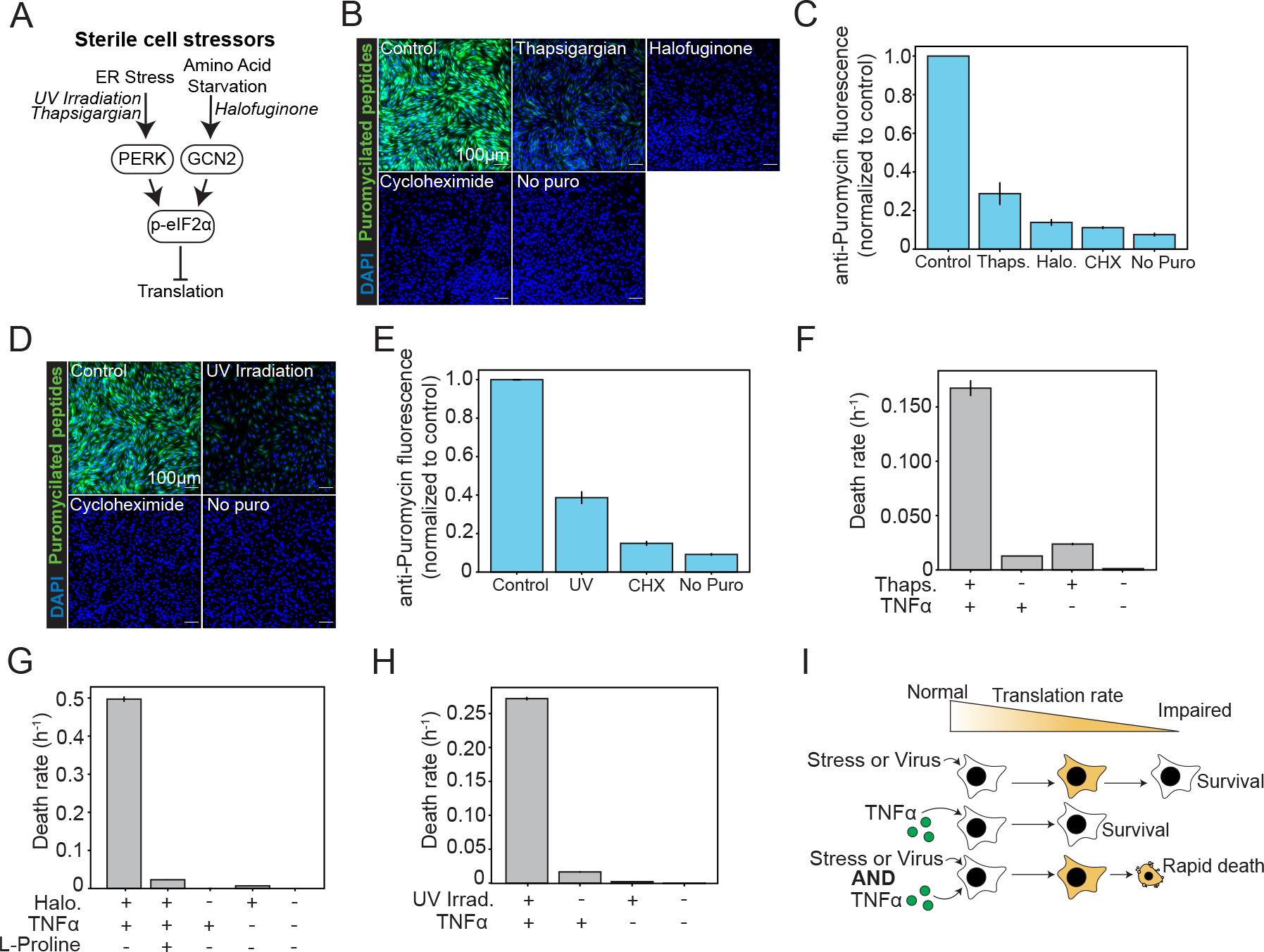
*Coupling cell death to TNFα safeguards cell viability during sterile perturbations to protein synthesis*. **(A)** Cartoon diagram showing examples of stressors that activate the ISR and drug inducers we use in our experiments. **(B)** Representative images of puromycin staining for cells treated with either 1 μM Thapsigargian or 500 nM Halofuginone for 6h prior to puromycin pulse. **(C)** Quantification of average puromycin fluorescence per cell for each of the different conditions. **(D)** Representative images of puromycin staining for cells that were UV irradiated with 150 J/m^2^. **(E)** Quantification of average puromycin fluorescence per cell for each of the different conditions. **(F-H)** Quantification of death rates from time lapse cell death experiments for each of the indicated conditions. For fig 6, in all gray bar plots, the error bars show the standard error of the fit. For all other plots the error bars show the mean ± s.d. of multiple biological replicates. **(I)** Cartoon depicting the working model. Both sterile stressors and viral infection can cause translational impairment in cells. However, in these cases, cells do not readily die. Low doses of TNFα also do not cause significant cell death. Combining translational inhibition due to stressors or viral infection causes rapid cell death.

## Discussion

In this study we examine how viral infection disrupts cellular proteostasis, and how cells guard for such disruptions and trigger immunity. We discovered that the extent of viral attenuation inversely correlates with translational inhibition: Infection with more virulent viruses leads to more inhibition. Consequently, productively-infected cells exhibit a significant and progressive decline in protein levels, and fail to trigger an antiviral response. The impact is particularly severe for rapidly degrading proteins, which decay quickly after global translational suppression. Specifically, halted translation leads to decay of the anti-apoptotic protein c-FLIP, which synergizes with TNFα to induce cell death and restrict viral spread. c-FLIP thereby guards for disruptions in protein synthesis by sensitizing cells to TNFα. Because TNFα is an inflammatory signal released by innate immune sentinels, it serves as a contextual cue of infection, enabling specificity in guard immunity during proteostatic stress.

### Productive infection impedes cellular activation of an antiviral response

Although the IFN response is central to antiviral immunity, our and others’ data show that viral replication is mutually incompatible with host translation of IFN-related proteins (Fig 1) ^1,34,47,71^. Although some host restriction factors are directly degraded by the virus, the majority of immune impairment is caused by global translational inhibition (Figs 1-2). Such translational inhibition is exquisitely and quantitatively tied to virulence. Consequently, infection with a highly-attenuated virus induces a canonical antiviral protein response, whereas infection with moderately-attenuated or wild type strains does not. The failure to upregulate immune proteins implies that productively-infected cells must rely on alternative forms of defense. We suggest that, *in vivo*, only cells that are abortively-infected, infected with DVGs, or uninfected cells neighboring them, will up-regulate immune proteins ^9^.

### Translational inhibition activates guard immunity

Guard immunity counters the rapid evolvability of pathogens by linking immunity to general signatures of virulence that disrupt cellular processes ^16^. Protein synthesis, which viruses absolutely require for replication, is one such example. Our study demonstrates that cells guard for disruptions in proteostasis via loss of c-FLIP, which decays rapidly upon cessation of translation (Figs 3 and 5). Reduced translation of c-FLIP synergizes with the cytotoxic cytokines TNFα and Fas to trigger cell death and limit viral spread (Figs 3 and 4). Our work also offers an explanation for the evolution of virus-encoded FLIP (v-FLIP) as a virulence factor in γ-herpesviruses and molluscipoxviruses ^72^. We expect viruses that encode v-FLIPs to evade this particular guard tactic.

Mathematical modeling provides clarity into the molecular logic that enables synergy between TNFα and translational inhibition: Through its’ action as a competitive inhibitor of Caspase 8 autocatalysis, c-FLIP modulates a stochastic molecular switch controlling induction of Caspase 8 positive feedback and triggering of cell death. Even a modest reduction in c-FLIP protein expression dramatically sensitizes cells to TNFα induced Caspase 8 activation, making cells die faster. Moreover, the stochastic noise present in the TNFα signaling network is instrumental for this process and confers an antiviral function.

### Conditioning cell death on TNFα safeguards viability in contexts of sterile stress

Various sterile environmental stressors perturb protein synthesis. Therefore, a major outstanding question was how cells distinguish sterile and nonsterile perturbations to protein synthesis, which is required to enact immunity. Our study provides an answer to this fundamental question by linking guard immunity to the presence of TNFα. Because TNFα and other cytotoxic cytokines are released by innate immune cells during infection, they contextualize translational stress as non-sterile, thereby enabling guard immunity. These observations reveal that guard immunity has evolved to work in concert with cytokines in higher organisms.

Our study suggests that the aberrant production of TNFα, or other cytotoxic cytokines, may have extremely deleterious effects on translationally-stressed cells in the absence of infection (Fig 6). We hope our work will spur the study of TNFα, cell stress, and translational inhibition in other physiologic and disease settings. Such settings may benefit therapeutically from TNFα blocking drugs or future efforts to ameliorate translational stress, or stabilize and enhance c-FLIP expression.

In conclusion, our work represents a major step forward in understanding mammalian guard immunity. Productive viral infection hamstrings the cells’ ability to induce antiviral proteins which, in synergy with TNFα, triggers guard immunity. This study demonstrates the interplay between innate immunity and guard immunity, and suggests that guard immunity mechanisms in vertebrates rely on innate immune signals to improve their fidelity.

## Methods

### Resource availability

Further information and requests should be directed to the lead contact, Dr. Jennifer Oyler-Yaniv and Dr. Alon Oyler-Yaniv

The raw data used to generate the figures in this paper are publicly-available from the Mendeley Data repository at the date of publication (Available upon publication).

All software used in this paper including for image analysis and modeling are available at our lab Github page: https://github.com/oylab

Any additional information required to reanalyze the data presented in this paper can be obtained from the lead contact upon request.

### Cell lines

Mouse NIH 3T3 cells and human A549 cells were purchased from ATCC (#CRL-1658; #CCL-185) and maintained in complete culture media: MEM supplemented with 10% heat inactivated fetal calf serum (Sigma, US Origin), 2mM L-glutamine, 100μg/ml of penicillin and 100μg/ml of streptomycin at 37°C and 5% CO_2_ in a humidified incubator.

### Viruses

HSV-1 strain KOS (wild type) and the related mutant strains d109 ^23^, 8ΔGFP ^24^, and dl38 were all provided by Dr. David Knipe (HMS), with permission from Dr. Neal DeLuca (University of Pittsburgh) for d109. HSV-1 strain KOS VP26-tdTomato or VP26-mCerulean were originally acquired from Dr. Prashant Desai (Johns Hopkins University) ^60^. Influenza A virus strain WSN 1933 was acquired from the BEI resource repository (#NR-3688).

### Viral propagation

To propagate wild type viruses, 4 × 10^6^ Vero cells were seeded in T150 cm^2^ tissue culture flasks, then infected with relevant strains at an MOI of 0.05. To infect, medium was removed and a 1.5 mL suspension of virus in PBS was added. After rocking briefly, the flask was transferred to 37°C and rocked every 10 min. After 1h, 45 mL of growth medium was added and incubated for 2 days at 37°C. After 2 days, cells were detached by shaking/tapping, and then pelleted for 20 minutes at 4°C. The supernatant was transferred to a conical tube and kept on ice. The remaining cell pellet was resuspended in 1 mL of growth medium, and cells were lysed by repeatedly pushing through a 1.5 mL syringe. 8 mL of growth medium was added to the lysate, and clarified by spinning for 10 minutes at 4°C. The two supernatant sources were then combined and centrifuged in a Beckman Coulter Optima L-100XP Ultracentrifuge at 40,000 g for 30 minutes at 4°C. After centrifugation, the supernatant was removed, and the remaining pellet was resuspended in MEM supplemented with 10% FBS and 10% glycerol to a final volume of 1.8 mL. The virus suspension was then homogenized using a 1.5 mL syringe and stored at -80°C. Viral titers were quantified by standard plaque assays performed in Vero cells.

### Imaging cell death or viral spread over time

2.5 × 10^4^ cells were seeded in 96 well glass bottom plate with high performance #1.5 cover glass (CellVis) pre-coated with 1-2.5 μg/cm^2^ fibronectin (Sigma). Cells were labeled with 20 ng/ml Hoechst 33342 (Thermo Fisher) and death was tracked by incorporation of 1:10,000 Sytox Green (Thermo Fisher).

### Cell lysate preparation for proteomics

4.0 × 10^6^ cells were seeded in 100 x 20 mm Corning Falcon Tissue Culture Dishes. After 24h, cells were infected with relevant viruses at an MOI of 10. At the specified time points, cells were washed with PBS and lifted with TrypLE for 3m. Cells were collected with 10 mL of PBS, spun at 200 g for 5 min, and washed 3x with 10 mL of ice cold PBS. The cell pellet was resuspended in 200 µL of proteomics lysis buffer composed of 6 M Guanidine Hydrochloride (Chem-Impex #29913), 2% Hexadecyltrimethylammonium Bromide (Millipore Sigma BioXtra #H9151), 100 mM Na+EPPS pH 8.0 (Millipore Sigma #E1894), 10 mM TCEP HCl (pH = 7.5) (Oakwood Chemical #M02624), and 1 roche tablet cOmplete (#50855100) per every 12.5 mL of lysis buffer.

### Sample Prep for LC-MS

Disulfide bonds were reduced by addition of 5 mM Dithiothreitol (DTT) (1 M stock in water) (Oakwood Chemical #3483-12-3) at 60°C for 15 min. Samples were sonicated for 10 rounds of 15 s at an amplitude of 60 on ice and clarified at 20k g for 15 min at 4°C (very minor to no pellet was observed). 50 mM of iodoacetamide (BioUltra, Millipore Sigma #I1149) (1 M stock in anhydrous N, N-Dimethylformamide (DMF (Millipore Sigma #227056))) was added to alkylate cysteines and samples were kept in the dark for 1 h. Following alkylation, samples were quenched by addition of 30 mM DTT. A chloroform/methanol protein precipitation was done as described previously ^73^. The protein disc was re-suspended at ∼2 µg/µL in 6 M GuHCl / 10 mM EPPS pH = 8.5. 50 µg of protein from each sample was diluted to 2 M GuHCl by addition of 10 mM EPPS pH = 8.5, and digested for 14 h with 20 ng/µL LysC (Wako). Samples were further diluted to 0.5 M GuHCl and an additional 20 ng/µL LysC was added along with 10 ng/µL Trypsin (Sequencing grade modified, Promega #V5111) and incubated at 37°C for 16 h. Solvent was removed in vacuo and each sample was re-suspended in 500 mM EPPS (pH = 8.0) at ∼1 µg/µL. 20 µL of TMTPro (Thermo, #A44520) (20 µg/µL in anhydrous Acetonitrile (ACN)) was added to 50 µg of peptides and incubated at room temperature for 2 h. 10 µL of 5% hydroxylamine (Millipore sigma #438227) was added to quench the reaction. Samples were combined and the solvent removed in vacuo. Samples were acidified to pH <= 1 with TFA, clarified at 20k g at 4°C for 10 min and desalted using a seppak (Waters). The seppak was wet with 1 mL of methanol, followed by 1 mL of 1% Formic acid (FA) followed by addition of sample. The seppak was washed twice with 1 mL of 1% FA and eluted into a 2 mL Eppendorf with 35% ACN/1% formic acid. Solvent from the eluent was removed in vacuo, the sample was re-suspended in 140 µL of 10 mM Ammonium Bicarbonate (pH = 8.0), and clarified in an ultracentrifuge at 70,000 rpm in a TLA-100 rotor for 30 minutes at 4°C. 100 µL of supernatant was fractionated by reverse phase chromatography as previously described ^74^ into a 96-well plate and 24 fractions were created, desalted by stage-tips ^75^ and ∼10 µg of peptides were subjected to LC-MS/MS analysis.

### LC-MS

A Thermo Orbitrap Eclipse was operated in data-dependent mode with a survey scan of 300-1100 m/z at an Orbitrap resolution of 120,000 with a 200 ms max injection time, AGC target of 1e6, and a Lens RF of 60%. The Eclipse was connected to a Thermo EASY-nLC 1200 HPLC. Individual runs were 4.5 hours in length with a gradient of 3% to 18% B (Solvent A 0.125% FA, 2% DMSO and Solvent B 0.125% FA, 80% ACN, 2% DMSO). Chromatography was performed with in house packed 100-360 µm inner-outer diameter microcapillary columns, which were manually packed with ∼0.5 – 1 cm of magic C4 resin followed by 40 cm of 1.8 cm roche resin ^76^. An in-house machined column heater was used and kept at 60°C. Peptides were ionized with a 2.6 kV voltage applied to a PEEK microtee. Four different experiments were used with different FAIMS ^77^ CVs (-32, -42, -52, -62). FAIMS was operated at standard resolution with a total carrier gas flow of 3.5 L/min. A dynamic exclusion window of 60 seconds with a filter of +/- 10 PPM was used. MS/MS isolations were performed with a quadrupole isolation window of 0.4 m/z, and peptides were fragmented with an HCD collision energy of 37%. The MS/MS scan was acquired with an orbitrap resolution of 50,000 with a first mass of 110 m/z. An AGC target of 50,000 was used with a maximum injection time of 120 ms. The low m/z TMT-Pro reporter ions were quantified with a 20 ppm mass tolerance and corrected for isotopic impurities as described previously ^78^.

### Data analysis - search

Data analysis and searches were performed essentially as described previously ^78^. The comet search engine was used instead of Sequest. A precursor peptide mass tolerance of 50 ppm was used allowing for the variable modifications of oxidation of methionine (+15.994914 Da) and deamidation of asparagine (+ 0.984016 Da). Iodoacetamide on cysteine (+ 57.02146372 Da) and TMT-Pro (+304.20660 Da) on n-termini and lysines were set as static modifications. The target-decoy approach was used to set the peptide false discovery at 1% ^79,80^ and these peptides were further filtered to obtain a 1% FDR on the protein level using protein pickr ^81^.

### Antiviral immune signature

The following GO terms and numbers were used to assemble a list of genes involved in the innate cellular response to virus and the type I IFNβ response: Antiviral Innate Immune Response, Regulation of Defense Response to Virus, Response to IFNβ, Cellular Response to IFN, Response to Virus (GO: 0140374, 0050688, 0035456, 0035458, 0009615, respectively). The following genes were added based on literature review: *Irf9, Ifi44, Tlr6, Oasl2, Ifit3b, Eif4ebp1, Xaf1, Igtp, Trim12c, Trim30a, Trim30d.* The mean expression of all genes in this list was used to generate the antiviral immune score. As an irrelevant gene set that is unrelated to antiviral signaling, we used genes listed in a growth factor signaling pathway or apoptosis signaling pathway with the following GO terms: MAPK cascade (GO: 0000165) Apoptotic signaling pathway (GO: 0097190).

### Immunofluorescence staining

Cells were fixed with 1.6% paraformaldehyde for 10 min on ice, then washed with PBS and permeabilized with ice-cold 90% methanol for at least 1 hour. Fixed and permeabilized cells were then washed 3x with PBS, and blocked (PBS, 0.3% Triton X-100, and 5% normal human serum (Jackson ImmunoResearch)) for 1h at room temperature. Note, human serum is important to block non-specific binding of full IgG to HSV-1 infected cells. In other instances, blocking buffer consists of PBS, 0.3% Triton X-100 and 3% Bovine Serum Albumin (Sigma). Cells were stained with antibodies for 30m-1h at room temperature or up to overnight at 4°C. Where, indicated, cells were then washed 3x with blocking buffer, then incubated with secondary antibodies and counterstained with DAPI for 1h at room temperature, light protected. Cells were then washed 3x with blocking buffer and an additional 2x with PBS before imaging.

### Puromycin incorporation assay

Cells were prepared with the indicated treatments, and then pulsed for 30min with 2 μM Puromycin (ThermoFisher) before fixing and permeabilizing ^49^.

### Antibodies

anti-ICP4 (clone 10F1, Abcam), anti-IAV Nucleoprotein (clone AA5H, Bio Rad), anti-Puromycin-Alexa 488 or 647 (clone 2A4, BioLegend) and anti-c-FLIP (clone D5J1E, Cell Signaling Technologies). All antibodies were used at 1:300 dilution of the stock. All polyclonal secondary antibodies were purchased from Jackson Immunoresearch and F(ab’)_2_ fragments were used at a concentration of 10 μg/ml. goat anti-Mouse IgG-Alexa 647 (cat: 115-606-003), goat anti-Mouse IgG-Alexa 594 (cat: 115-586-003).

### Microscopy

Images were collected on a confocal Nikon Eclipse Ti2E-inverted microscope equipped with a Nikon CFI Plan Apo λ 20×, numerical aperture (NA) 0.75 objective lens. Images were acquired with a CrestOptics X-Light V3 light engine and a Photometrics Kinetix camera. For live-cell imaging, cells were maintained at 37°C and 5% CO_2_ in a humidified environmental chamber. All imaging was accomplished using custom automated software written using Python and Micro-Manager.

### Treatment with cytokines, drugs and small molecules

Cells were treated with the following concentrations of molecules, unless otherwise indicated in the text. Mouse TNFα (5 ng/ml, Peprotech), Human TNFα (5 ng/ml, Peprotech), cycloheximide (CHX) (20 μM, Sigma), anisomycin (ANM) (5 μM, Sigma), bortezomib (2.5 μM, Cayman chemical), carfilzomib (2 μM, Dr. Marc Kirschner, HMS), Z-VAD(Ome)-FMK (30 μM, Cayman chemical), Necrostatin-1 (30 μM, Cayman chemical), anti-CD95 agonist antibody (1 μg/ml, clone JO2, BD Biosciences), Obatoclax (10 μM, Cayman chemical), A1210477 (10 μM, Cayman chemical), LCL-161 (5 μM, Cayman chemical), Thapsigargian (1 μM, Cayman chemical), Halofuginone (500 nM, Dr. Ralph Mazitschek, Mass General Hospital), L-Proline (4 mM, Sigma).

### UV Irradiation

The media was removed from cells, then they were washed once with PBS and kept in PBS with plastic cover removed during irradiation for 15s with a UV lamp set to 10 Joule/m^2^/s. After irradiation, complete media was replaced.

### Transfections and siRNA knockdown

For transfection of poly dA-dT, HSV-1 *Vhs*, and SARS-CoV-2 *Nsp1*, 2.5 × 10^4^ cells were seeded in 96 well glass bottom plate and adhered overnight. 200 ng/well of plasmids or poly dA-dT were transfected with Lipofectamine 3000 according to manufacturer protocol.

For siRNA knockdowns, cells were sequentially knocked down. First, 6 well plates were seeded at a density of 1.25 × 10^5^ per well and cells were adhered overnight. Cells were then transfected with either a negative control (scramble) siRNA (Thermo Fisher #AM46111) or a pooled combination of three *Cflar*-targeting siRNAS (Thermo Fisher #’s 66164, 66073, and 65977) targeting exons 2, 3, and 5. siRNAS were all transfected at a final concentration of 10 nM using Lipofectamine 3000 in absence of P3000 reagent, as specified by the manufacturer instructions. After 24 h, cells were trypsinized and re-seeded at 2.5 × 10^4^ cells per well in 96 well plates and simultaneously transfected again with the same final concentrations of siRNAs. Experiments were then initiated 24 hours after the second transfection.

### Western blotting

Cells were harvested in NuPAGE LDS Sample Buffer (Invitrogen) for 10 min at 70°C and sonicated to shear genomic DNA. Proteins were separated by electrophoresis using 4-12% Bis-Tris mini protein gels and transferred to PVDF membranes (BioRad). Blots were incubated with primary antibodies at 4°C and then with HRP conjugated secondary antibodies for 1 hour at room temperature. HRP was detected using ECL substrates (Thermo Scientific) and Licor Odyssey Fc Imager.

### Image analysis

Automated image analysis was accomplished using custom software written in Python. Image analysis software is available from our GitHub repository: https://github.com/oylab/oyLabImaging

We segmented individual cell nuclei using a seeded watershed or Stardist ^82^ algorithm applied to the nuclear dye (Hoechst 33342). Incorporation of a nucleic acid stain (Sytox green) was used to call cell death based on an increase in the relevant fluorescence intensity. The threshold for death was determined by comparing to uninfected or untreated control conditions where cell death is minimal. We used the same strategy to call infected cells based on the fluorescence intensity of VP26-mCerulean or -tdTomato HSV-1 strains. To determine the fraction of infected or dead cells, we divided the number of dead cells at each timepoint by the total number of cells (alive or dead) identified by segmentation at the indicated frame.

### Mathematical modeling

We write a stochastic differential equation that incorporates TNFα triggered Procaspase-8 cleavage, autocatalysis of Procaspase-8 by Caspase-8 dimers, inhibition of Caspase-8 by free c-FLIP, natural Caspase-8 degradation, and stochastic biochemical noise. TNFα signaling is modeled as a Hill function with coefficient 1 and EC_50_ of 1 ng/ml. Noise is modeled as a simple additive Gaussian process. We use a quasi-steady state approximation to couple the levels of free c-FLIP to the current levels of Caspase-8.

To arrive at a population wide measurement of survival, we used the python package py-pde to solve the associated Fokker-Planck equation, which describes the time evolution of the probability density function of Caspase-8 activity, and measured the fraction of live cells in the population as a function of time. The initial conditions were chosen to be the steady state distribution under conditions of maximal c-FLIP translation and no TNFα signaling, as is the case in healthy cells. The fraction of dead cells was calculated as the total probability of having Caspase-8 activity greater than 50% of its maximal value as a function of time. Death rates were extracted by fitting these curves to a simple exponential fit. The code to execute the model is available here (Available upon publication).

### Statistical Analysis and Data Presentation

All relevant data are shown as mean ± standard deviation (s.d.), or mean ± 95% confidence intervals of relevant fit. Statistical tests were selected based on appropriate assumptions with respect to data distributions and variance characteristics.

## Author Contributions

Conceptualization (JO-Y, AO-Y, MS), Investigation (JO-Y, AO-Y, MS, EC, SB, RT, MB), Software (AO-Y, MS), Writing (JO-Y, AO-Y, MS), and Resources (JO-Y, LP, MS)

## Acknowledgements

We thank all members of the OY lab for critically reading the manuscript, Roy Wollman (UCLA) and Tim Mitchison (HMS) for helpful conversations, Talley Lambert at the Center for Imaging Technology Education (HMS) for assisting with development of our image analysis software, Rui Tong (HMS) for providing plasmids, Caroline Mock (HMS) for facilitating UV irradiation, David Knipe (HMS), Neal DeLuca (University of Pittsburgh), and Prashant Desai (John Hopkins University) for viruses, and Ralph Mazitschek (MGH), Ying Lu (HMS), and Marc Kirschner (HMS) for reagents. MS is supported by a grant from the Dean of Harvard Medical School. MB is supported by the Swiss National Science Foundation postdoc mobility fellowship.

## Declaration of Interests

The authors declare no competing interests.

---

**Supplementary Figure 1:**
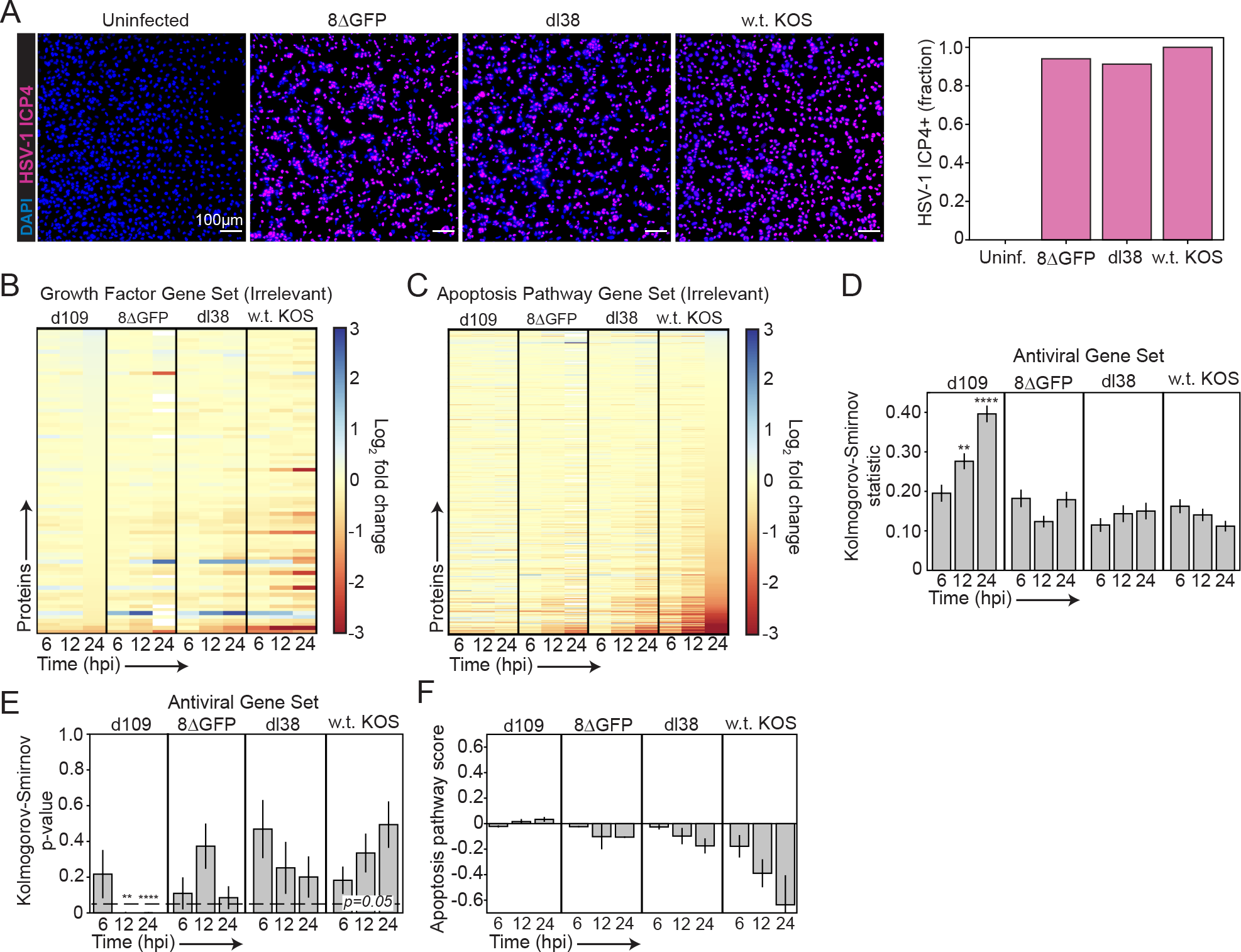
**(A)** Representative images and quantification of HSV-1 infected cells (based on ICP4 immunofluorescence) after overnight infection with MOI 10 of each of the different viral mutants. **(B-C)** Heatmaps showing protein expression for an irrelevant set of genes associated with MAPK signaling (B) or apoptosis signaling (C). **(D)** Quantification of the Kolomgorov-Smirnov statistic for all proteins listed in our antiviral immune signature compared to 10,000 randomly-selected genes **(E)** Quantification of the Kolomgorov-Smirnov p-value for the same conditions as in D. **(F)** Quantification of the apoptosis protein expression score for cells infected with each of the viral mutants. To compute the score, we took the mean expression of all proteins annotated in the GO apoptosis pathway list ± s.e.m. for each protein.

**Supplementary Figure 2:**
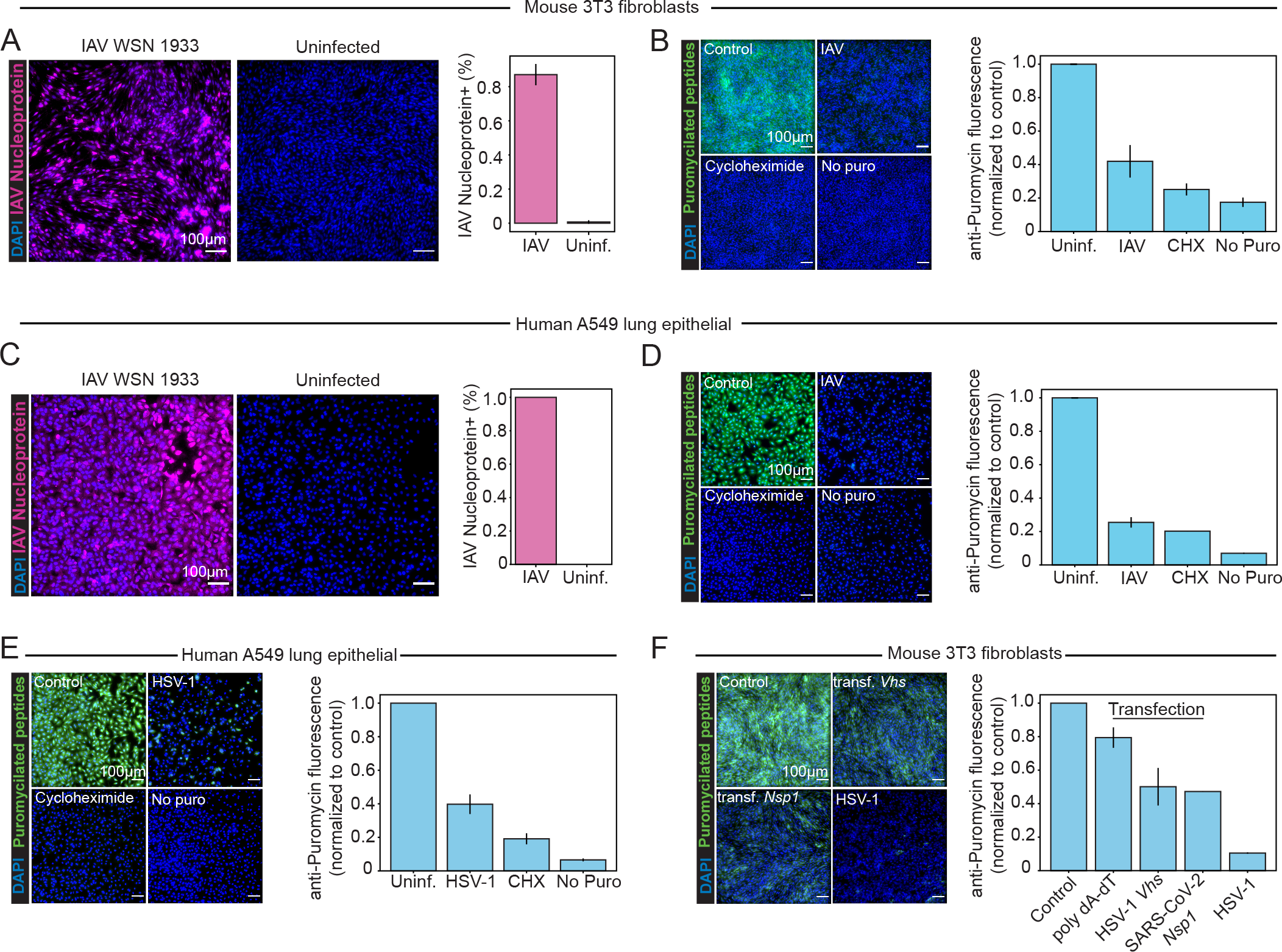
**(A)** Representative images and quantification of IAV infection based on Nucleoprotein (NP) immunofluorescence after overnight infection of 3T3 fibroblasts with MOI 10. **(B)** Representative images and quantification of puromycin staining for fibroblasts infected overnight with MOI 10 of IAV. **(C)** Representative images and quantification of IAV infection (based on NP immunofluorescence) after overnight infection of A549 cells with MOI 10. **(D)** Representative images and quantification of puromycin staining for A549 cells infected overnight with MOI 10 of IAV. **(E)** Representative images and quantification of puromycin staining for A549 cells infected overnight with MOI 10 of HSV-1. **(F)** Representative images and quantification of puromycin staining for fibroblasts transfected with 200 ng/well of plasmids encoding either HSV-1 *Vhs*, or SARS-CoV-2 *Nsp1.* All bar plots show the mean ± s.d. of multiple biological replicates.

**Supplementary Figure 3:**
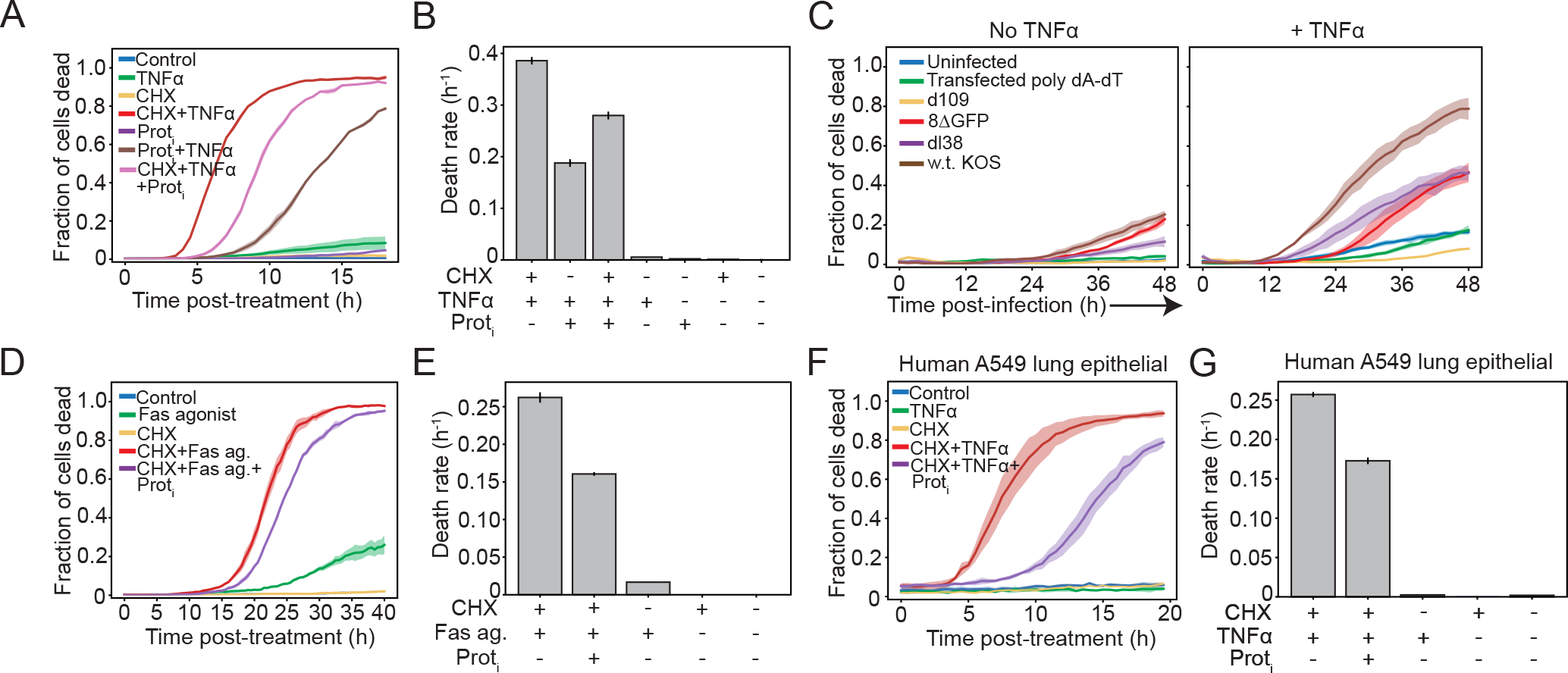
**(A)** Quantification of cell death over time for cells treated with either TNFα, CHX, or our Prot_I_ alone, or a combination of TNFα and CHX, or TNFα and Prot_I_. **(B)** Quantification of death rates fit from A. **(C)** Quantification of cell death over time for cells infected with MOI 10 of each viral strain, or transfected with 200 ng/well of poly dA-dT with or without concurrent treatment with TNFα. Accompanies Fig 3I. **(D)** Quantification of cell death over time for cells treated with either 1 μg/ml anti-CD95 (Fas agonist), anti-CD95 with CHX, or anti-CD95 with CHX and Prot_I_. **(E)** Death rates fit from D. **(F)** Quantification of cell death over time for A549 cells treated with either 5 ng/ml TNFα or CHX, or a combination of TNFα and CHX, or TNFα and CHX and ProtI. E **(G)** Death rates fit from F. For all bar graphs in this figure, the error bars show the standard error of the fit. For all other plots, the data show the mean ± s.d. of multiple biological replicates.

**Supplementary Figure 4:**
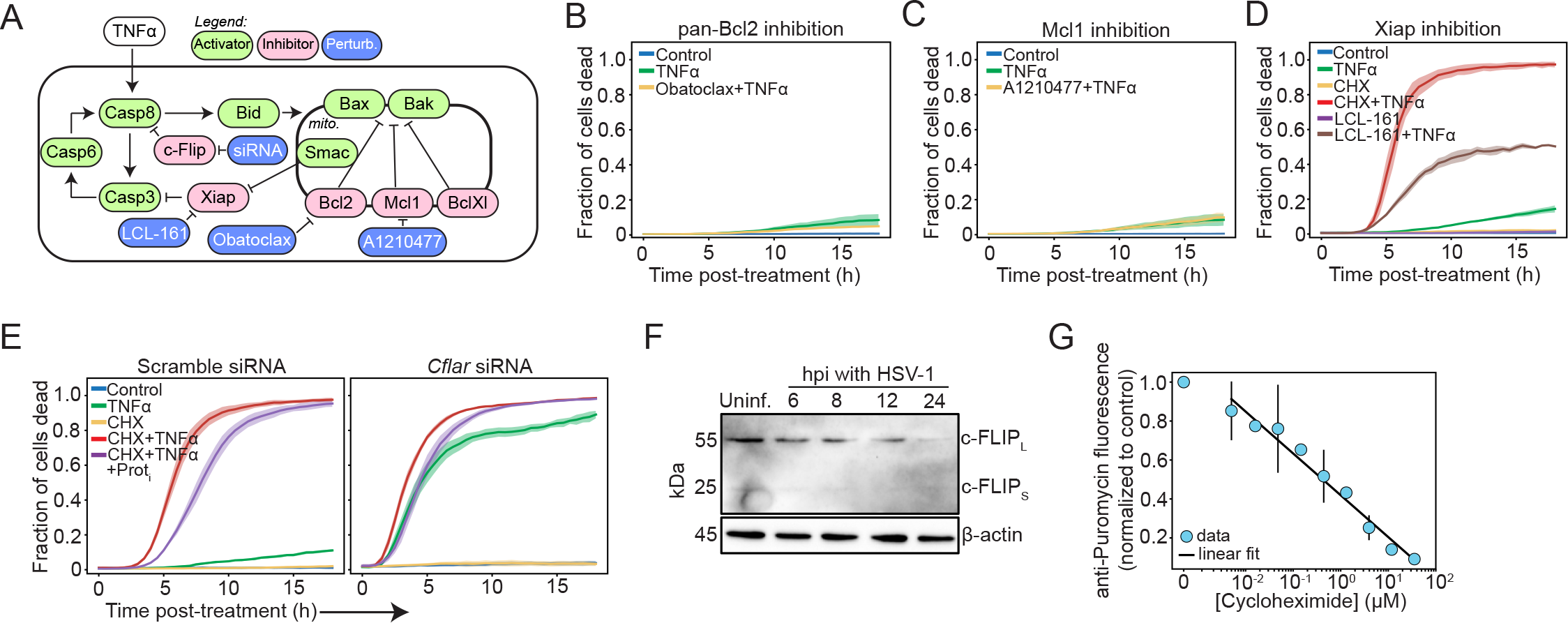
**(A)** Cartoon diagram of the key positive and negative regulators of the extrinsic apoptotic signaling network, and various perturbation strategies. **(B)** Quantification of cell death over time for cells treated with either TNFα alone or TNFα combined with 10 μM of the pan-Bcl2 inhibitor Obatoclax. **(C)** Quantification of cell death over time for cells treated with either TNFα alone or TNFα combined with 10 μM of the Mcl1 inhibitor A1210477. **(D)** Quantification of cell death over time for cells treated with the indicated treatments. LCL-161 was used at a concentration of 5 μM. **(E)** Quantification of cell death over time for cells treated with control or *cflar* siRNAs and the indicated conditions. Accompanies Fig 5B-C. **(F)** Western blot for mouse c-FLIP after infection with w.t. HSV-1. **(G)** Quantification of puromycin incorporation for cells treated with the indicated concentration of cycloheximide, and fit linearly. For figure S4, all data show the mean ± s.d. of multiple biological replicates.

**Supplementary Figure 5:**
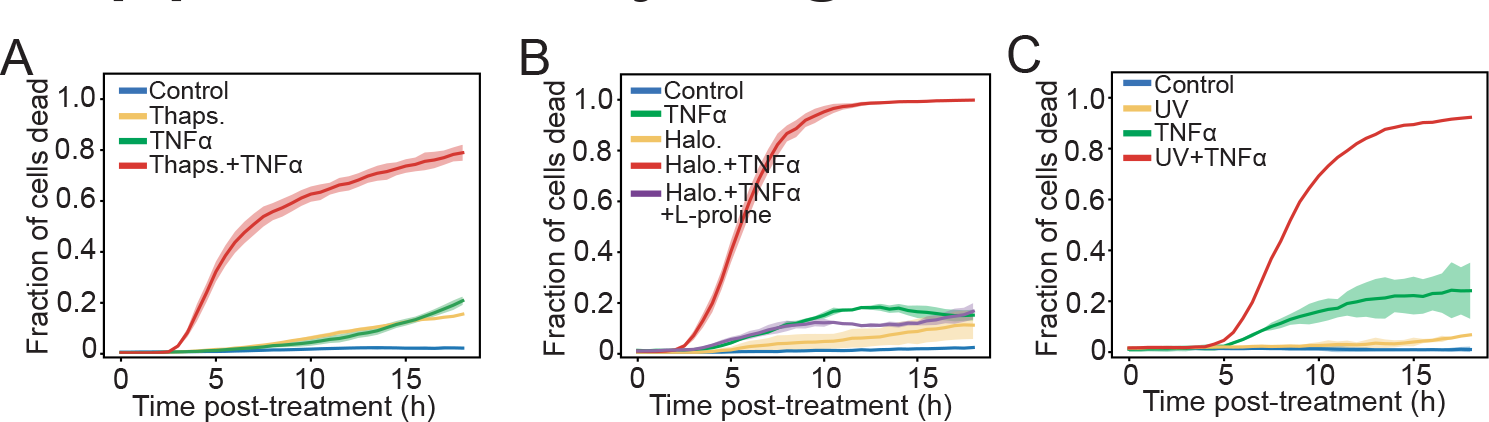
**(A)** Quantification of cell death over time for cells treated with TNFα or Thapsigargian, or TNFα and Thapsigargian. Accompanies Fig 6F. **(B)** Quantification of cell death over time for cells treated with TNFα or Halofuginone, or TNFα and Halofuginone, or TNFα and Halofuginone and 4mM L-Proline. Proline competes for the target of Halofuginone (prolyl tRNA synthetase). Hence, proline elimination of cytotoxicity indicates that cytotoxicity arises from Halofuginone’s on-target mechanism. Accompanies Fig 6G. **(C)** Quantification of cell death over time for cells treated with TNFα or UV irradiation, or TNFα and UV irradiation. Accompanies Fig 6H. For Fig S5, all of the data show the mean ± s.d. of multiple biological replicates.

## Notes

### Competing Interest Statement

The authors have declared no competing interest.

